# Theory of Mind and Discourse Production in Schizotypy: An fMRI Study

**DOI:** 10.1101/2025.09.19.677472

**Authors:** Sharlene D. Newman, Madeleine Rein, Emily Gann, Yanyu Xiong

## Abstract

**Background:** Schizotypy (ST) reflects subclinical traits linked to schizophrenia spectrum disorders and associated cognitive and social impairments. Theory of Mind (ToM) and discourse production deficits are well-documented in schizophrenia (SZ), yet the neural basis of discourse-related ToM processes in ST remains unclear. This study investigated brain activation during narrative planning and production in individuals with schizotypal traits.

**Methods:** Thirty young adults (mean age = 18.8 years) completed standardized assessments, including the Schizotypal Personality Questionnaire–Brief Revised (SPQ-BR), adverse childhood experiences (ACEs), depression (PHQ-9), and dissociation (DES-B). Participants performed a discourse task in an fMRI scanner, describing nine-panel cartoons requiring inference of character intentions. Behavioral discourse metrics included total and inferred events. fMRI analyses examined activation during planning and production phases, with SPQ-BR positive, negative, and disorganized traits entered as regressors.

**Results:** Schizotypal traits correlated with multiple psychosocial risk factors, including elevated depression, ACEs, and dissociation (r = .48–.82, p < .01). During planning, canonical ToM/self-referential regions (vmPFC, precuneus, insula) were recruited. Positive traits correlated with increased activation in the right temporo-parietal junction, precuneus, and lingual gyrus, whereas disorganized traits were associated with reduced activation in the precuneus and lingual gyri. During production, networks spanning vmPFC, hippocampus, right TPJ, and basal ganglia were engaged. Negative traits correlated with increased motor/premotor activation, while disorganized traits correlated with reduced activation in lingual gyrus, SMA, and cerebellum.

**Conclusions:** Findings demonstrate distinct neural correlates of schizotypal traits during discourse planning and production, supporting models of schizophrenia-spectrum risk emphasizing disrupted inference and integration processes.

## Introduction

Schizophrenia (SZ) is a debilitating psychiatric disorder in which patients may lose touch with reality and experience a constellation of positive, negative, and disorganized symptoms, including hallucinations, delusions, alogia, avolition, flat affect, and disorganized speech (WHO, 2022). Affecting approximately 24 million people, SZ is among the leading causes of disability worldwide (WHO, 2022). Recent research conceptualizes SZ not as a discrete category but as part of a psychosis spectrum, with symptoms ranging from normative functioning to full clinical presentation (Grant et al., 2018). Within this spectrum, schizotypy (ST) represents a constellation of latent personality traits resembling subclinical manifestations of SZ, including magical thinking, unusual perceptual experiences, and reduced speech output (Cohen et al., 2015). ST is of particular importance as it may serve as a risk factor for the development of SZ and related spectrum disorders. Moreover, individuals with high ST demonstrate functional impairments akin to those observed in SZ, including reduced interpersonal connectedness, occupational and educational difficulties, and financial strain (Cohen et al., 2015; Haas et al., 2015). Cognitive studies also reveal overlapping deficits, such as impaired verbal and visuospatial working memory (Siddi et al., 2017). Because schizotypal traits are expressed in non-clinical populations without the confounds of antipsychotic medications or hospitalization histories, ST provides a valuable model for probing the biological and environmental mechanisms underlying SZ.

One key domain disrupted in SZ, and potentially in ST, is Theory of Mind (ToM), the ability to infer others’ beliefs, emotions, and intentions (Vucurovic et al., 2021, 2023). ToM lies at the intersection of several higher-order cognitive processes, including mentalization, social cognition, and pragmatic language (Schurz & Perner, 2015; Haas et al., 2015). Communication of ToM constructs requires discourse production, and deficits in both ToM and discourse are consistently observed in SZ (Parola et al., 2018). For instance, up to 30% of individuals with SZ present with co-occurring pragmatic, cognitive, and social impairments (Bambini et al., 2016). However, there is a fundamental difference in the ToM processes used during common mentalizing task paradigms and the mentalizing processes occurring during speech production (Frith, 2004). Despite this, our current understanding of impairments in ToM in SZ populations is heavily based on task paradigms that require little to no discourse production. Therefore, the present study addresses this gap by using a novel discourse production task to capture mentalizing processes occurring during speech production.

At the neural level, ToM tasks typically elicit activation in the temporo-parietal junction (TPJ), medial prefrontal cortex (MePFC), and right inferior frontal gyrus (rIFG) (Carrington & Bailey, 2009). Discourse production recruits overlapping fronto-temporal networks (Mason and Just, 2006, 2009). SZ patients consistently demonstrate impairments across these domains, with fluctuating severity depending on clinical state (Frith & Corcoran, 1996; Drury et al., 1998; Sarfati, 1999). Deficits in metacognition are thought to contribute to impaired social functioning and emotional understanding (Lysaker, 2014). Disorganized cognition, even when controlling for verbal IQ, is associated with reduced semantic processing efficiency (Haas et al., 2015; Tonelli, 2014). During speech tasks, SZ patients produce less relevant and more tangential discourse, marked by weak associations and unclear referential connections (Marini et al., 2008; Haas et al., 2015). Some evidence suggests that persecutory delusions may exacerbate ToM deficits (Langdon et al., 2005), though others argue these impairments reflect broader dysfunction observed across psychiatric conditions (Chan et al., 2025). Neuroimaging studies implicate differential activation in the dorsomedial and orbitofrontal cortices, inferior prefrontal cortex, superior temporal sulcus, and amygdala during affective ToM processing in SZ (Caillaud et al., 2020). Importantly, ToM impairments are observed not only in patients after a first psychotic episode but also in first-degree relatives and high-risk populations (Bora & Pantelis, 2013; Nelson et al., 2008), underscoring their potential role as an endophenotype.

One of the remaining questions is whether the differential TOM related brain activity is also observable in individuals with ST. Although schizotypy research is more limited, emerging evidence suggests parallel deficits. Affective ToM studies in ST report altered activation in regions such as the left medial temporal gyrus (MTG), parahippocampal gyrus, lingual and fusiform gyri, and atypical patterns of medial prefrontal and temporo-parietal junction activity (Kronbichler et al., 2017). Behavioral work also finds subtle language disturbances, such as loosened associations and logical inconsistencies, thought to stem from working memory deficits (Kiang et al., 2007). Early adulthood may represent a particularly sensitive window for detecting these alterations, as it coincides with both heightened schizotypal expression and the typical onset period for SZ (Tonini, 2010). Despite these findings, few studies have directly examined the neural correlates of discourse production in ST populations, leaving the relationship between ToM, language, and brain function incompletely understood.

The present study addresses this gap by investigating neural activation associated with discourse production in individuals with schizotypal traits. Participants completed a narrative task requiring them to describe nine-panel cartoons, which necessitated both inference of character intentions (ToM) and coherent language production. This paradigm provides an opportunity to assess whether ToM-related discourse engages distinct or altered neural networks in ST, thereby offering insight into mechanisms of social communication deficits and potential risk pathways for schizophrenia spectrum disorders. Due to the lack of prior literature assessing ToM processes from speech production, we implemented a novel manual annotation protocol to capture “visual” and “inferred” events (see methods) previously used in our prior research assessing ToM in high schizotypy from a college student sample (Gann et al., 2025). We anticipated the relationship between discourse-level ToM and ST to be modulated by activation in neural networks associated with ToM.

## Methods

### Participants

Twenty-nine individuals participated in the study (6 male; aged 18.77±.73 years). This study was conducted in accordance with the ethical principles of the Declaration of Helsinki and was approved by the Institutional Review Board at The University of Alabama. Informed consent was obtained from all participants prior to their involvement in the study.

### Assessments

Participants were recruited from a pool of individuals who participated in an online study. That study used Qualtrics to obtain demographic information (e.g., age, gender, race) and the following assessments:

- Schizotypal Personality Questionnaire-Brief Revised (SPQ-BR; Cohen et al., 2010). The SPQ-BR contains 32 questions in which participants responded on a 5-point Likert scale (strongly disagree to strongly agree) assessing positive, negative, and disorganized schizotypal traits. The SPQ-BR has demonstrated to be replicable and valid for undergraduate samples (Davidson et al., 2016).
- The Adverse Childhood Experiences Questionnaire (ACEs; Felitti et al., 1998) was administered. It is a 10-item measure that quantifies adverse or traumatic experiences (e.g., physical, emotional, and sexual abuse) that occurred before the age of 18.
- Depressive symptoms were assessed using the PHQ-2 (Spitzer et al., 1999), a two-item screener with a sensitivity of 79% and a specificity of 86% for any depressive disorder (Lowe et al., 2005). Anxiety was assessed using the two-item GAD-2 (Spitzer et al., 2006)
- The Alcohol Use Disorders Identification Test-C (AUDIT-C; Bush et al., 1998) is a 3-item screening tool to assess alcohol consumption, drinking behaviors, and alcohol-related problems. Scores range from 0 to 12. A score of 4 or more for men and 3 or more for women indicates hazardous drinking and indicate possible alcohol use disorder.
- The Cannabis Use Disorders Identification Test-Revised (CUDIT-R; Adamson et al., 2010) is an 8-item screening measure valid for identifying likely cases of DSM-5 cannabis use disorder (CUD). Scores range from 0 to 32 and are categorized as safe, hazardous, or disordered. A score of 8 or more indicates hazardous use, and 12 or more indicates possible CUD.
- The Brief Dissociative Experiences Scale (DES-B) - Modified (Dalenberg & Carlson, 2012) is an 8-item measure assessing the severity of dissociative experiences. Each item is rated on a five-point scale for the past 7 days, and the total score can range from 0 to 32. The average score was used for the analysis. It reduces the overall score to a 5-point scale, which allows researchers to think of the severity of an individual’s dissociative experiences on the 0-4 scale: 0-none, 1-mild, 2-moderate, 3-severe, or 4-extreme4. The use of the average total score was found to be reliable, easy to use, and clinically useful in the DSM-5 Field Trials.

Participants from the on-line survey were invited to participate in the current study based on their SPQ-BR score and were asked to complete an in-person assessment battery that included:

- Substance use was evaluated with the Alcohol, Smoking, and Substance Involvement Screening Test (ASSIST). The ASSIST is an 8-item questionnaire that evaluates the lifetime and current use of various psychoactive substances as well as any problems associated with substance use (Humeniuk et al., 2010).
- Depressive symptoms were evaluated using the Patient Health Questionnaire-9 (PHQ-9). The PHQ-9 is a 9-item Likert scale questionnaire (not at all to nearly every day) that has been shown to be a reliable and valid evaluation of the severity of depressive symptoms (Kroenke et al., 2001).
- Participants were then administered the MATRICS Consensus Cognitive Battery (MCCB). The MCCB is an assessment battery used to evaluate global cognitive functioning in those with schizophrenia. It has been shown to be reliable and valid in schizophrenic populations as well as older adolescent populations (Nuechterlein, 2008). Participants were administered 8 component tests that examined speed of processing (Trail Making Test (TMT); Category Fluency; Animal Naming; Brief Assessment of Cognition in Schizophrenia-Symbol Coding (BACS-SC)), verbal learning (Hopkins Verbal Learning Test-Revised; HVLT-R), non-verbal working memory (Weschler Memory Scale-Third Edition (WMS-III): Spatial Span), verbal working memory (Letter-Number Span (LNS)), visual learning (BVMT-R), and social cognition (Mayer-Salovey-Caruso Emotional Intelligence Test (MSCEIT): Managing Emotions).
- Finally, participants completed The Awareness of Social Inference Test (TASIT) Part 2: Social Inference-Minimal (McDonald et al., 2003). The TASIT examines social understanding and cognition through 15 short video clips that show varying social situations. Participants are questioned on the characters’ feelings, thoughts, intentions, and meanings of their words. It has been validated in schizophrenic and adolescent populations (McDonald et al., 2017; McDonald et al., 2006).

### fMRI Task

Participants completed a discourse production task in an MRI scanner. They were asked to verbally describe a story that matched a nine-panel cartoon (see Figure 1) on a coil mirror that reflected the image displayed on a screen at the back of the scanner. The comics depicted a cat in various scenarios. The comics were presented for 60 seconds. The participants were asked to review all panels and develop the story and then press a button when they were ready to begin speaking. When they finished speaking, they pressed a button to indicate they completed the description. There were 10 comic panels divided into 2 blocks. There was a break between the blocks. Stimuli were presented using PsychPy 2023.2 on a 2017 iMac. Speech was recorded using the MR compatible FOMRI-III + NC Opto-acoustic microphone.

**Figure 1:**
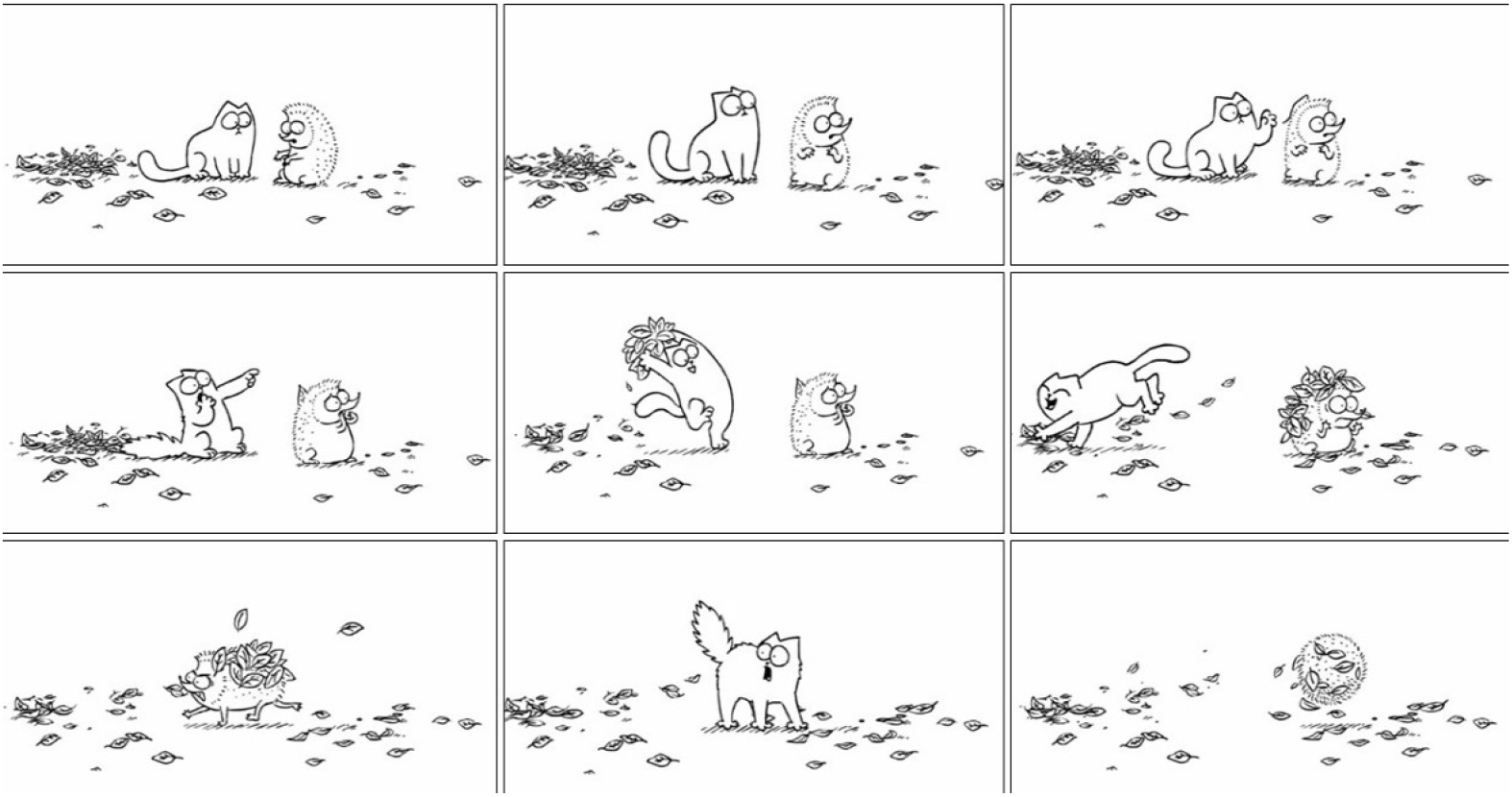
An example of a 9-panel comic used as stimuli for fMRI discourse task. Cartoons are from Simon’s Cat (https://www.simonscat.com/).

Speech recordings were manually transcribed. Annotators were trained on the coding scheme developed. Raters were instructed to annotate for visual and inferred events. Visual events were defined as an action that could be explicitly seen occurring in the comic (e.g., The cat ran away) while inferred events were defined as instances where the speaker described an event that was not explicitly presented such as inferring a character’s thoughts, feelings, or (e.g., He was angry). Raters were blind to subject demographics and SPQ-BR scores during discourse analysis. The metrics were total number of events (sum of visual and inferred events), and proportion of inferred events (visual events divided by total number of events). Degree of mentalizing was operationalized as proportion of inferred events. Discrepancies were discussed. The interrater reliability was 0.82.

### fMRI acquisition

Participants were scanned with a Siemens 3-T MAGNETOM Prisma MRI scanner at the MRI facility of the University of Alabama using the software syngo MR XA30 and a 32-channel head coil. An echo planar Imaging sequence with free induction decay was used to acquire functional data with the multi-band acceleration factor equal to 4. Each volume consisted of 72 slices with 2mm thickness (TR = 2s; TE = 0.03s; Flip Angle = 52°; isotropic voxel size = 2 × 2 × 2mm^3^; field of views [FOV] = 216mm). A whole-brain high resolution T_1_-weighted structural image was acquired using magnetization prepared rapid gradient echo sequence (MPRAGE) accelerated with GRAPPA parallel imaging. Each volume has 208 slices of 1mm thickness (TR = 2.3s; TE = 0.003s; Flip Angle = 9°; isotropic voxel size = 1 × 1 × 1mm^3^; FOV = 256mm).

### fMRI data analysis

MRI data were processed and analyzed using SPM12 [University College London; (Ashburner, 2012)]. The preprocessing steps included: slice timing correction, motion correction using a rigid body realignment algorithm, co-registration, spatial normalization using the MNI template in SPM12 (IXI549space) and each participant’s T1 scan, and smoothing with the Gaussian kernel filter of 8 mm. The final voxel size after normalization was 2 × 2 × 2 mm^3^. The amount of head motion was closely examined and no subject showed excessive translational movements > 1 mm. FIRMM (Turning Medical) was used to monitor motion during scanning.

Event-related responses were analyzed using a general linear model (GLM) with the planning and production phases and 6 motion estimates as regressors. The second-level random-effect analysis was performed by entering the first level contrast of each subject of each run in a regression analysis that included the three measures from the SPQ-BR – positive, negative and disorganized traits. Family-wise correction with a p<0.05 and an extent threshold of 50 voxels was used to examine the activation related to the planning and production phases. Activation correlated with each SPQ-BR measure was also explored using a critical threshold of *p* uncorrected <.005 with cluster size 100 was used. The activation locations were determined using xjView toolbox (https://www.alivelearn.net/xjview).

## Results

### Behavioral results

Summary statistics are presented in Table 1. A correlation analysis was performed between the assessments and SPQ-BR scores. Verbal Learning was negatively correlated with both positive (r=-0.43, p=0.02) and negative symptoms (r=-0.5, p=0.0054). PHQ-9 was positively correlated with disorganized (r=0.48, p=0.0087), positive (r=0.56, p=0.0016) and negative symptoms (r=0.62, p=0.0003). ACEs score was correlated with disorganized (r=0.59, p=0.0014), positive (r=0.68, p=0.0001) and negative symptoms (r=0.53, p=0.0055). Dissociation was correlated with disorganized (r=0.82, p<0.0001), positive (r=0.81, p<0.0001) and negative symptoms (r=0.78, p<0.0001). SSAR was not significantly related to any symptoms (r=-0.35, p=0.059).

**Table 1:**
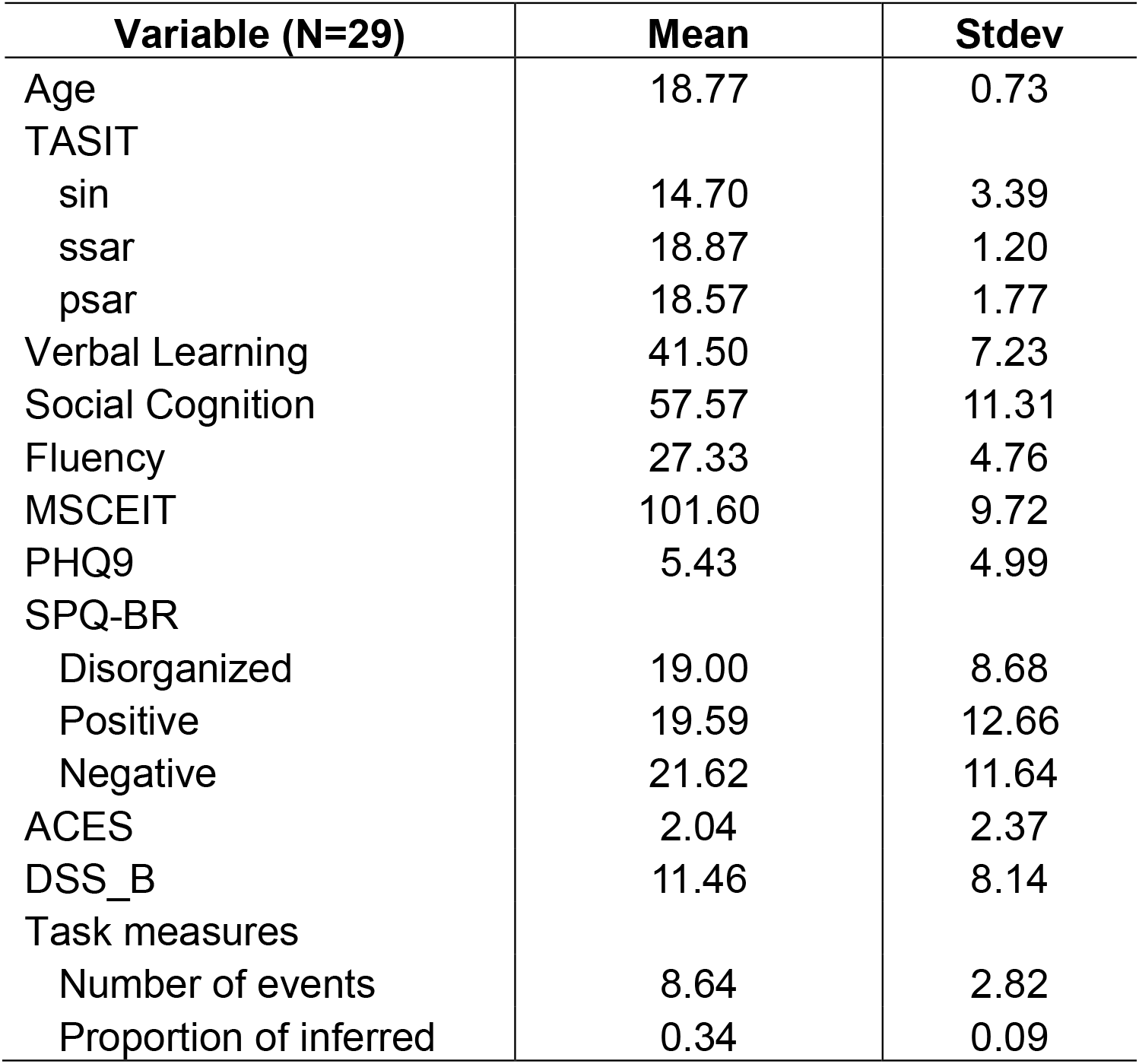
Sample characteristics.

#### Planning

Twenty-seven participants’ data were included in the MRI analysis (two were excluded due to poor data quality). Activation during the planning phase (threshold FWE, p<0.05, extent 50 voxels) was observed in the ventromedial prefrontal cortex, posterior cingulate/precuneus and the right insula (see Figure 2 and Table 2). The three sub-factors of the SPQ-BR were entered into the model as regressors (threshold uncorrected p<0.005, extent 100 voxels). Positive traits were positively correlated with the right TPJ, precuneus cortex, and lingual gyrus. Disorganized traits were negatively correlated the precuneus and bilateral lingual gyri. Negative traits failed to elicit significant correlation with brain activation.

**Table 2:** Planning Activation.

| Table 2: Planning Activation |  |  |  |  |  |  |
| --- | --- | --- | --- | --- | --- | --- |
| region |  | k | T | x | y | z |
| Right Precuneus |  | 129 | 10.5 | 24 | -44 | 14 |
|  |  |  | 8.48 | 16 | -38 | 18 |
| Left Precuneus |  | 114 | 8.23 | -8 | -70 | 28 |
|  |  |  | 6.79 | -8 | -58 | 22 |
| Left Precuneus |  | 67 | 7.54 | -20 | -46 | 12 |
|  |  |  | 6.67 | -14 | -38 | 16 |
| Left Posterior Cingulate |  | 122 | 7.49 | 0 | -30 | 44 |
|  |  |  | 6.77 | 0 | -20 | 34 |
| Right Insula |  | 58 | 7.49 | 38 | 8 | 10 |
|  |  |  | 7.23 | 46 | 2 | 8 |
| Ventromedial Prefrontal Cortex |  | 51 | 6.74 | 0 | 46 | 2 |
| Positive Traits |  |  |  |  |  |  |
| region |  | k | r | x | y | z |
| Left Precuneus |  | 278 | 0.68 | -4 | -66 | 18 |
| Right Temporo-parietal Junction |  | 125 | 0.65 | 66 | -36 | 24 |
| Lingual Grys |  | 109 | 0.64 | 0 | -90 | 0 |
| Disorganized Traits |  |  |  |  |  |  |
| region |  | k | r | x | y | z |
| Left Lingual Gryus |  | 288 | -0.69 | -12 | -52 | 4 |
|  |  |  | -0.63 | -6 | -68 | -2 |
|  |  |  | -0.53 | -12 | -78 | 6 |
| Right Lingual |  | 130 | -0.66 | 2 | -92 | 0 |
| Left Precuneus |  | 258 | -0.64 | -12 | -76 | 32 |

**Figure 2:**
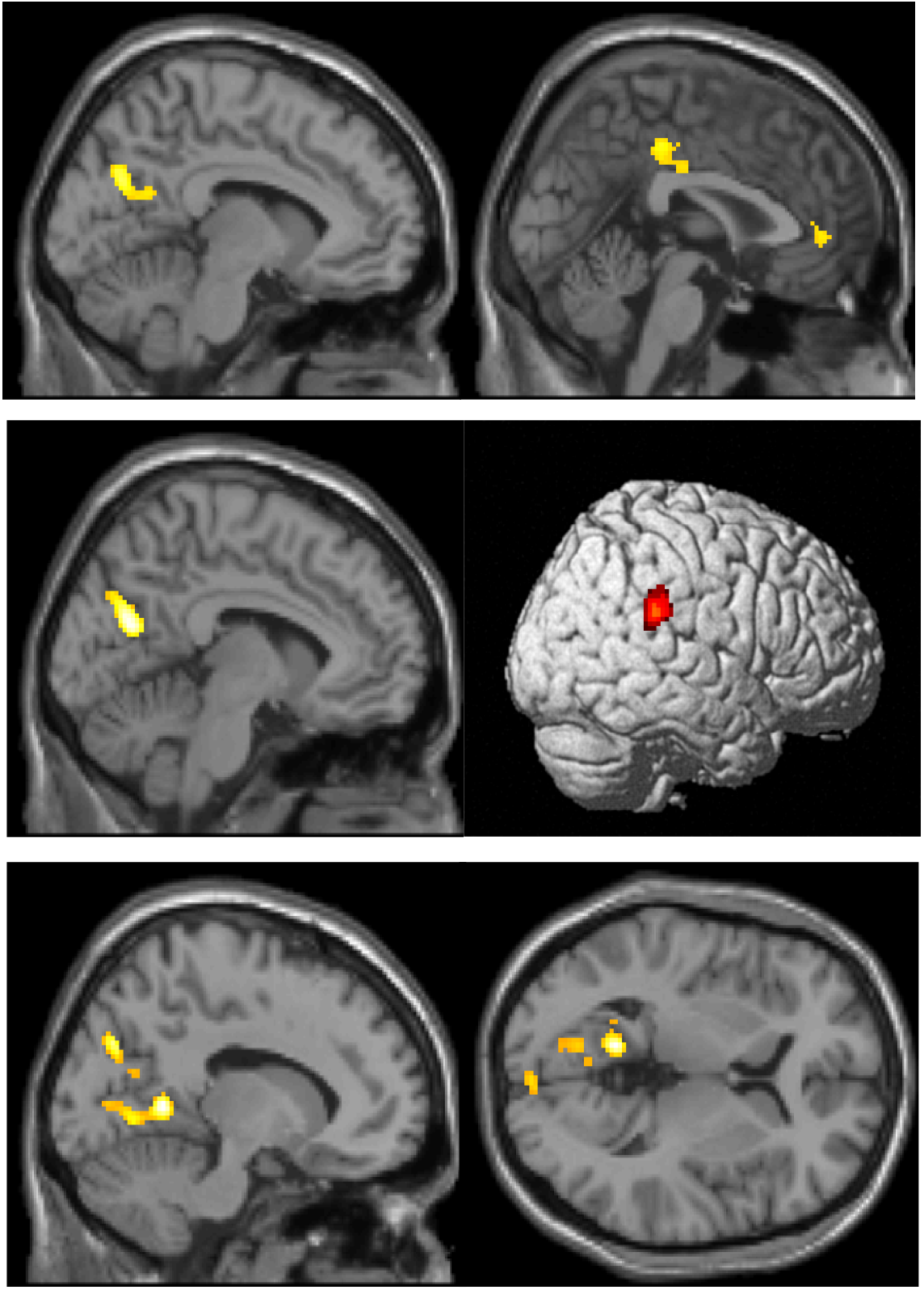
Top: activation related to planning. Middle: positive correlation with positive traits. Bottom: negative correlation with disorganized traits.

#### Production

Activation during the production phase (threshold FWE, p<0.05, extent 50 voxels) was widespread and included regions active during the planning phase (the ventromedial prefrontal cortex, precuneus, posterior cingulate) as well as the right TPJ, basal ganglia (caudate nucleus, putamen), hippocampus and middle frontal gyrus (see Table 3 and Figure 3). Negative traits were positively correlated with motor and pre-motor cortex bilaterally, and the supplementary motor area (SMA). Disorganized traits were negatively correlated with the lingual gyrus, middle cingulate/SMA, and the anterior lobe of the cerebellum. Positive traits failed to elicit significant correlation with activation.

**Table 3:**
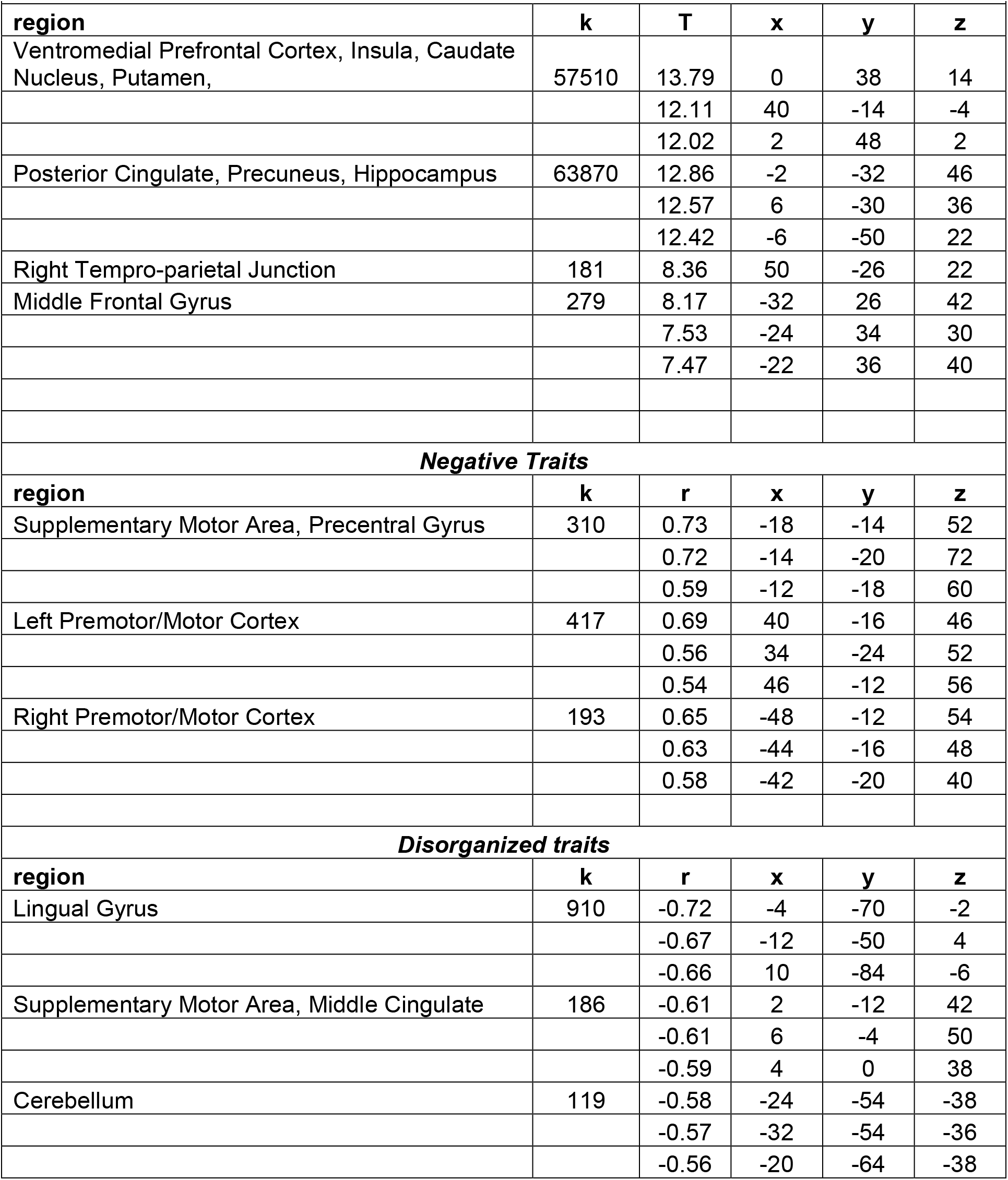
Production Activation.

| **Table 3: Production Activation** |  |  |  |  |  |
| --- | --- | --- | --- | --- | --- |
| **region** | **k** | **T** | **x** | **y** | **z** |
| Ventromedial Prefrontal Cortex, Insula, Caudate Nucleus, Putamen, | 57510 | 13.79 | 0 | 38 | 14 |
|  |  | 12.11 | 40 | -14 | -4 |
|  |  | 12.02 | 2 | 48 | 2 |
| Posterior Cingulate, Precuneus, Hippocampus | 63870 | 12.86 | -2 | -32 | 46 |
|  |  | 12.57 | 6 | -30 | 36 |
|  |  | 12.42 | -6 | -50 | 22 |
| Right Temporo-parietal Junction | 181 | 8.36 | 50 | -26 | 22 |
| Middle Frontal Gyrus | 279 | 8.17 | -32 | 26 | 42 |
|  |  | 7.53 | -24 | 34 | 30 |
|  |  | 7.47 | -22 | 36 | 40 |
| **Negative Traits** |  |  |  |  |  |
| **region** | **k** | **r** | **x** | **y** | **z** |
| Supplementary Motor Area, Precentral Gyrus | 310 | 0.73 | -18 | -14 | 52 |
|  |  | 0.72 | -14 | -20 | 72 |
|  |  | 0.59 | -12 | -18 | 60 |
| Left Premotor/Motor Cortex | 417 | 0.69 | 40 | -16 | 46 |
|  |  | 0.56 | 34 | -24 | 52 |
|  |  | 0.54 | 46 | -12 | 56 |
| Right Premotor/Motor Cortex | 193 | 0.65 | -48 | -12 | 54 |
|  |  | 0.63 | -44 | -16 | 48 |
|  |  | 0.58 | -42 | -20 | 40 |
| **Disorganized traits** |  |  |  |  |  |
| **region** | **k** | **r** | **x** | **y** | **z** |
| Lingual Gyrus | 910 | -0.72 | -4 | -70 | -2 |
|  |  | -0.67 | -12 | -50 | 4 |
|  |  | -0.66 | 10 | -84 | -6 |
| Supplementary Motor Area, Middle Cingulate | 186 | -0.61 | 2 | -12 | 42 |
|  |  | -0.61 | 6 | -4 | 50 |
|  |  | -0.59 | 4 | 0 | 38 |
| Cerebellum | 119 | -0.58 | -24 | -54 | -38 |
|  |  | -0.57 | -32 | -54 | -36 |
|  |  | -0.56 | -20 | -64 | -38 |

**Figure 3:**
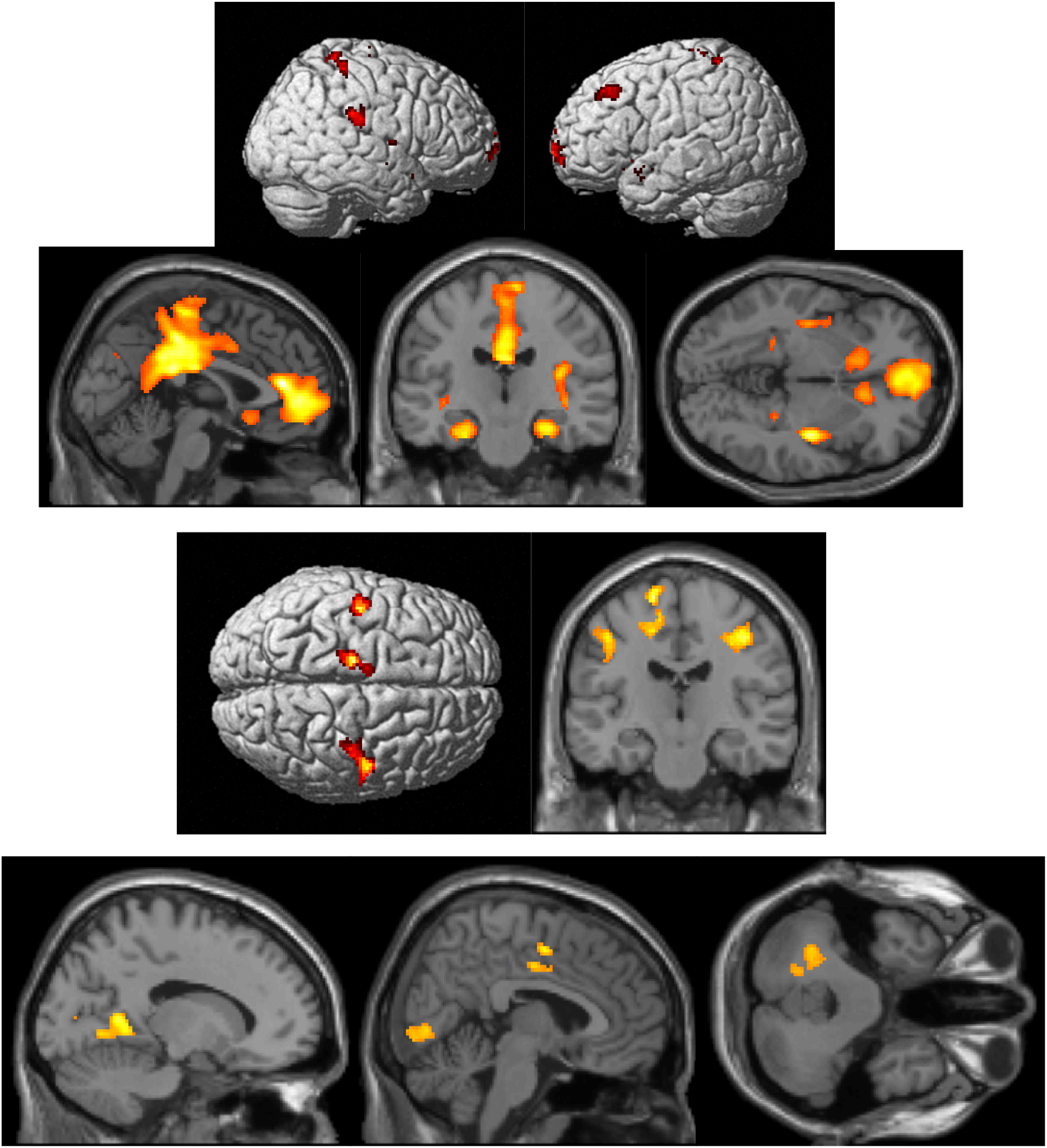
Top: production activation. Middle: positive correlation with negative traits. Bottom: negative correction with disorganized traits.

## Discussion

The present study examined discourse production and neural activation in relation to schizotypal traits in a non-clinical sample. Consistent with prior work, the correlation analysis showed that schizotypy was associated with impairments in verbal learning (Cohen et al., 2015; Siddi et al., 2017), elevated depressive symptoms (Webster et al., 2022), and greater exposure to adverse childhood experiences (Toutountzidis et al., 2022). These findings reinforce the broader literature indicating that schizotypy confers functional impairments across cognitive and affective domains and may represent an important risk phenotype for schizophrenia spectrum disorders. Although we did not observe significant correlations between schizotypal traits and behavioral discourse measures, fMRI data revealed distinct associations between schizotypy dimensions and brain activation in regions implicated in Theory of Mind (ToM), language, and social cognition. These findings extend prior evidence that schizotypal traits affect discourse-related neural networks even in the absence of overt speech abnormalities.

### Planning Phase: Situation Model Construction

During the planning phase, participants recruited canonical ToM and self-referential processing regions including the ventromedial prefrontal cortex (vmPFC), posterior cingulate cortex/precuneus, and right insula. These regions are consistent with networks previously implicated in the construction of situation models and in the anticipation of others’ mental states (Frith & Corcoran, 1996; Schurz & Perner, 2015). Importantly, positive schizotypal traits were correlated with activation in the right temporo-parietal junction (TPJ), lingual gyrus, and precuneus, regions central to inferring intentions, monitoring social agents, and constructing mental perspectives. This suggests that individuals with elevated positive traits may engage ToM-related regions more heavily during narrative planning, possibly reflecting compensatory strategies or altered salience attribution.

In contrast, disorganized traits were negatively correlated with activation in the precuneus and bilateral lingual gyri. Given that the precuneus plays a critical role in perspective-taking and integrating multimodal information, hypoactivation may underlie difficulties in generating coherent situation models, consistent with discourse disturbances observed in schizotypy and schizophrenia (Marini et al., 2008; Haas et al., 2015).

### Production Phase: Language, Motor, and Memory Integration

The production phase engaged a broader network including the vmPFC, precuneus/posterior cingulate, hippocampus, right TPJ, basal ganglia, and middle frontal gyrus, consistent with the integration of memory, speech production, and ToM processes. Negative schizotypal traits were positively correlated with activation in motor and premotor cortices, as well as the supplementary motor area (SMA). This finding aligns with reports that negative symptoms in schizophrenia are linked to psychomotor slowing and speech production deficits (Vöckel et al., 2023). Increased recruitment of motor regions may reflect compensatory effort to generate fluent speech in the context of diminished motivational and expressive resources.

Disorganized traits were associated with reduced activation in the lingual gyrus, SMA/middle cingulate, and cerebellum. These regions are implicated in perceptual integration, motor sequencing, and error monitoring. Reduced engagement may contribute to fragmented or tangential discourse by impairing the smooth integration of sensory, motor, and cognitive signals.

### Interpreting Disorganized Traits: A Circular Inference Framework

The strong association between disorganized traits and altered perception-related activation can be interpreted through the lens of circular inference models. These frameworks propose that schizophrenia-spectrum pathology involves disrupted excitatory/inhibitory (E/I) balance, leading to reverberations between top-down priors and bottom-up sensory evidence (Bishop, 2007; Friston, 2008; Markov et al., 2013; Leptourgos et al., 2017). Such disruptions foster “seeing what we expect” and “expecting what we see,” producing perceptual distortions and incoherent inferences (Jardri et al., 2017). Our findings suggest that even in non-clinical populations, disorganized schizotypal traits are linked to reduced precision in perceptual integration, which may manifest behaviorally as incoherent discourse or difficulty organizing thought (Kwapil et al., 2008; Krężołek et al., 2019).

### Limitations and Future Directions

Several limitations should be acknowledged. The small sample size (N=30) limits statistical power and generalizability. The behavioral discourse measures may not have been sufficiently sensitive to capture subtle differences, and future studies could incorporate more granular linguistic analyses (e.g., semantic coherence, acoustic markers). Longitudinal designs are needed to establish whether the observed neural correlates predict functional decline or psychosis onset. Finally, integration with electrophysiological measures (e.g., ERPs indexing early sensory processing) may help clarify the temporal dynamics of circular inference and E/I imbalance.

## Conclusion

In summary, this study demonstrates that schizotypal traits are differentially associated with neural activation in ToM, perceptual, and motor networks during discourse planning and production. Positive traits were linked to increased activation in ToM-related regions, negative traits to motor recruitment, and disorganized traits to reduced perceptual and integration processes. These findings align with models of schizophrenia-spectrum disorders emphasizing disrupted inference and integration mechanisms and highlight the value of studying schizotypy as a window into early psychosis risk.

## Notes

### Competing Interest Statement

The authors have declared no competing interest.

### Summary of Updates

There were significant errors in the references and additions that were needed.

